# Simultaneous Detection of Dopamine and Serotonin with Carbon-based Electrodes

**DOI:** 10.1101/2021.08.31.458352

**Authors:** Gaurang Khot, Neil Shirtcliffe, Tansu Celikel

## Abstract

Graphite-based materials, like pyrolyzed carbon electrodes, are widely used as implantable electrochemical sensors, for the detection of neurotransmitters, neuromodulators, and gaseous species, thanks to their strong mechanical properties, superior electron-transfer kinetics, and in-vivo stability. Electrochemical properties of graphite can be improved by coating them with carbon nanotubes (CNTs) which improves sensitivity, selectivity, and resistance to biofouling. Although several types of electrodes have been developed to detect biologically relevant targets like monoamines, multiplexed sensing of dopamine and serotonin is not yet widely available. Herein we introduce pyrolyzed carbon electrodes coated with CNTs for fast scan cyclic voltammetry for simultaneous detection of dopamine and serotonin with a sensitivity of 52±8nA/μM and 5±17nA/μM, respectively. Serotonin shows a broad oxidation peak at 0.68V±0.12V. When dopamine and serotonin are probed simultaneously at 10 Hz, dopamine oxidizes at 0.1V± 0.1 and serotonin oxidizes at 0.78V±0.12 and dopamine reduces at −0.35V±0.1 and serotonin at 0.1V±0.2 V. Thus the sensors shows discrimination between dopamine and serotonin and are suitable for simultaneous detection of these monoamines.

## Introduction

Carbon based materials have received widespread attention as neuronal implants because of their stable electrochemical kinetics, biostability and bioinert nature (Aragona et al., 2009; Khot et al., 2021; Venton et al., 2002). These properties of carbon based electrodes are widely exploited to study the neurochemical communication of the brain for understanding sensorimotor control, learning, reward, and motivation (Hughes and Celikel, 2019; Robinson et al., 2003; Stagkourakis et al., 2019). Graphite based materials are considered as workhorses for neuro-electrochemistry for detection of neurotransmitters and neuromodulators, because of their sensitivity, reduced potentials and stability compared to metal electrodes (Zachek et al., 2008). Considerable interests exists in increasing the sensitivity and stability of graphite electrodes by performing chemical modifications by means of anionic membrane and electrochemical pretreatment for improving sensitivity(Heien et al., 2003; Ross and Venton, 2012). While these methods have allowed increase in sensitivity, drawbacks of electrochemical pretreatment is the adsorption of cationic molecules leading to biofouling of graphite (Bennet et al., 2013; Jacobs et al., 2014; Swamy and Venton, 2007). Another drawback of graphite based electrodes is the disintegration of the electrodes during prolong implantations, thus leading to immune reactions from nearby brain regions (Bennet et al., 2013, 2016).

Carbon nanotubes (CNTs) are increasingly used as an alternative electron carrier material because of their ability to resist biofouling, stability and superior electrochemical kinetics compared with graphite electrodes (Mendoza et al., 2020; Schmidt et al., 2013; Swamy and Venton, 2007). Their long term stability and reduced potentials as compared to graphite based materials have seen the application of CNT based materials in form of CNT yarns (Mendoza et al., 2020; Yang et al., 2016, 2017) and blending of CNTs onto suitable matrix materials (Alba et al., 2015; Taylor et al., 2019) for electrochemical detection of neurotransmitters (Schmidt et al., 2013)

Serotonin is a monoamine neurotransmitter that originates from raphe nuclei in the brain (Adell et al., 2002; Schubert et al., 2015). Serotonergic neurons in the brain mature early during embryonic development and critically involved in the functional development of the brain and behavior, including controlling mood, eating, sleeping, (Karayol et al., 2021), and learning (Azarfar et al., 2019; Homberg, 2012). Low levels of serotonin in the brain are known to cause depression (Lacasse and Leo, 2005) which impacts decision making, learning, and attention (Homberg, 2012). Given the widespread role played by serotonin in developmental, neurological, cognitive, and pathophysiological conditions (Azarfar et al., 2019; Homberg, 2012) it is important to develop sensors that can detect serotonin levels. Challenges associated with electrochemical detection of serotonin are low basal concentration of serotonin, biofouling of the probes and rapid uptake thus limiting the availability time in extracellular domain (Hashemi et al., 2012).

Traditionally electrochemical detection of serotonin is done using “Dopamine waveform” (Heien et al., 2003), wherein a triangular waveform is cycled from from −0.4V to 1.3V at 400V s^-1^ at 10 Hz. Application of this waveform results in the adsorption of serotonin on the surface of the electrodes, which results in biofouling of probes, thereby limiting the shelf life of the electrode (Hashemi et al., 2009). To address this problem a modified waveform was introduced by restricting the anodic switching potential 1.0V and limiting the cathodic potential at −0.2V at 1000V s^-1^ to limit the adsorption of serotonin on the surface of the electrodes (Jackson et al., 1995). These changes allowed prolonged measurements, improved sensitivity and selectivity for serotonin (Hashemi et al., 2009, 2011, 2012). The application of specific waveform allows sensitive detection of serotonin, however limits the multiplex detection of dopamine and serotonin (Hashemi et al., 2009). Previous attempts on multimodal detection of dopamine and serotonin using graphite electrodes coated with CNTs shows the oxidation of monoamines at common oxidation window and their resolution is done using reduction peaks (Swamy and Venton, 2007). This approach limits the identification of analytes in a complex matrix like the brain, thus requiring a need for material that is able to perform multiplex detection of dopamine and serotonin, without causing biofouling of electrodes from electrochemical side product 5-Hydroxyindoleacetic acid (5-HIAA) whose basal concentration is 10 folds higher than serotonin (Hashemi et al., 2009; Jackson et al., 1995). To overcome these limitations and simultaneously measure dopamine and serotonin in the same waveform, herein we introduce pyrolytic electrodes coated with CNTs for multiplexed measurement of monoamines.

## Materials and Methods

All chemicals, dopamine hydrochloride (DA-HCl), serotonin hydrochloride (5-HT-HCl) ascorbic acid, 4,-(2-hydroxyethyl-)1-piperazineethanesulfonic acid (HEPES), single-walled carbon nanotubes (SWCNTs), sodium chloride (NaCl), potassium chloride (KCl), sodium bicarbonate (NaHCO_3_), magnesium chloride (MgCl_2_), monosodium phosphate (NaH_2_PO_4_), and 5-hydroxyindoleacetic acid (5-HIAA), were purchased from Sigma Aldrich. Quartz capillaries (o.d. 1 mm, i.d. 0.5mm, length 7.5 cm were purchased from Sutter Instruments.

### 1. Preparation of Carbonized Electrodes

Quartz capillaries (O.D: 1.0mm and I.D.: 0.5mm) were pulled to a submicron diameter of 0.8-1.5μm using a Sutter puller p-2000 (Sutter instruments). The pulling parameters were, heating temperature 750°C, the velocity of 50, and the delay of 127. The tip was broken shorter to reach a final diameter of 25-30μm. Once the desired diameter was achieved pyrolysis was performed using Propane gas at 1 barr against counter flow of 50mL min-^1^ nitrogen. Once the tip was carbonized to the desired extent, they were allowed to be cooled in a nitrogen environment. For electrochemical detection, electrodes were coated with CNT solution suspended in Nafion (Khot et al., 2021).

### 2. Electrochemical setup

Electrochemical detection of serotonin was carried out using a patch-clamp amplifier (Intan Technologies, Los Angeles, USA). A silver wire coated in KCl solution (3.5M) was used as a reference electrode and a pyrolytic-CNT coated electrode was used as a working electrode. The electrochemical chamber allowed mimicking of transient changes in concentrations that were identical to that of electrophysiological recordings (Kole et al., 2020; da Silva Lantyer et al., 2018). To mimic the transient changes of neurotransmitters in the brain, a buffer was delivered to the recording chamber at a constant flow rate of 2mL min^-1^ and the analyte was introduced as a bolus on 5 s, followed by 10 s pause (Schmidt et al., 2013). Electrochemical detection was carried out by using “Serotonin waveform” (Hashemi et al., 2009; Jackson et al., 1995) with modifications, 0.2V to +1.3V and cycled back to −0.4V at a scan rate of 1000V s^-1^ at 10Hz. For electrochemical measurements of neurotransmitters, quartz capillaries were heated for 60 s, dipped in CNTs dispersed in Nafion solution (5 mg mL^-1^ in ethanol). Electrochemical measurements were carried in artificial cerebrospinal fluid (aCSF) as described before (Kole, 2015; Kole et al., 2017b, 2017a, 2018) and included 135mM NaCl, 5.4mM KCl, 5mM Na-HEPES buffer, 1.8mM CaCl_2_ and 1mM of MgCl_2_.

### 3. Computational Models

To visualize the interactions between CNTs and serotonin, we constructed a ball and stick model in Avogadro (Hanwell et al., 2012) and their molecular interactions were calculated in ORCA (Neese, 2012, 2018). CNTs model consisted of a small chain of CNTs (2,1) where the ends and sides were capped with hydrogen or oxygen. Geometry optimization of CNTs was done using the Density Functional Theory (DFT) calculated in ORCA. To understand the binding of serotonin onto the CNT surface, first serotonin’s geometry was optimized using implicit solvent mode in ORCA. To facilitate the transition search, serotonin and CNTs were connected by means of dummy atoms, and no bond or distance constraints were laid. The structures were optimized in ORCA and the resultant output file was loaded in Avogadro for visualization.

## Results and Discussions

### 1. Probing Serotonin Dynamics

To investigate the electrochemical kinetics of serotonin onto graphite electrodes, we introduced 1*μ*M of serotonin in the flow chamber, using the “Dopamine waveform” (Heien et al., 2003) at a scan speed of 400V s^-1^. The waveform allowed us to detect serotonin, wherein serotonin oxidized at 0.7V (±0.1V) and reduced at −0.1V (**Figure 1A**). While the waveform was selective, we observed biofouling of electrodes after 4 injections of serotonin. To address the issue of biofouling, we coated carbonized probes with CNTs and applied “Dopamine Waveform” at 600 V s^-1^ (**Figure 1B**). While we could observe a change in electrochemical kinetics of serotonin, this waveform was susceptible to biofouling as well. Using CNT-coated electrodes, we saw a broad oxidation peak at 0.65 ±0.18V and a reduction peak at 0.2 V±0.1V. To address the issue of biofouling and enable long-term measurements, we made use of the “Serotonin waveform” (Hashemi et al., 2009; Jackson et al., 1995). Through an iterative process, we found that extending the anodic waveform to 1.3V and the cathodic scan to −0.4V improves sensitivity for detection of serotonin. Application of this waveform at 600 V s^-1^ shows an oxidation peak at 0.72V(±0.05V) and a secondary oxidation peak is seen at 0.45V, a reduction peak can be seen at 1.1V, which we suspect occurs from deprotonation of serotonin-indole group complex and reduction peak seen at −0.3V is from the reduction of serotonin (**Figure 1C**). To study the electrochemical kinetics, the scan rate was plotted against peak current (**Figure 1D**), thus suggesting that serotonin is transported from bulk to the surface of the electrode by adsorption. We then converted the current, redox signatures against the number of trials in a hotspot (**Figure 1E**) and we see that two distinct oxidation states can be seen from 0.7V to 1.3V while reduction can be seen from 0.0V t0 −0.4V. This likely suggests that the oxidation of serotonin on pyrolytic-CNT electrodes is multistep.

**Figure 1:**
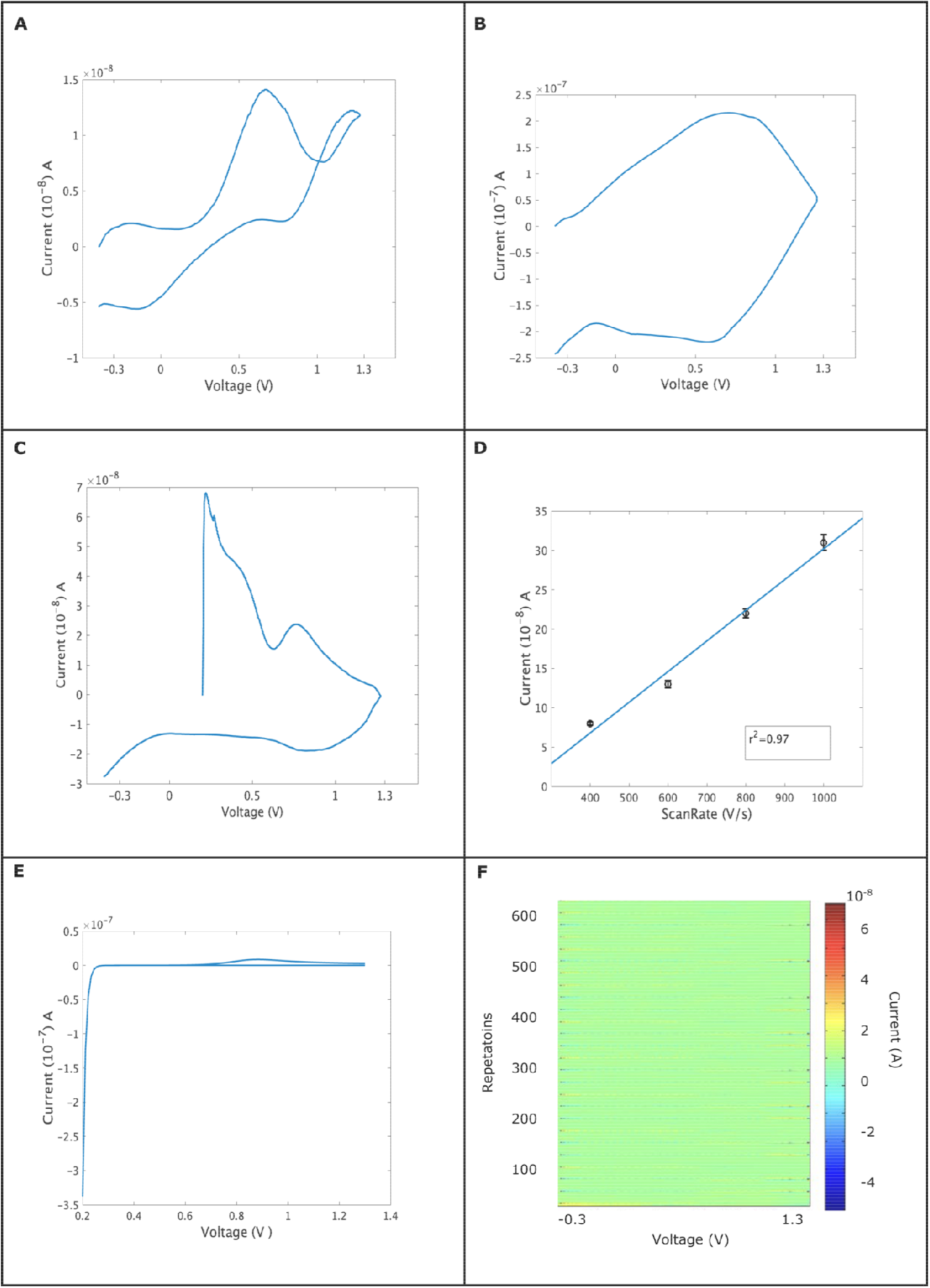
Shows the electrochemical kinetics for 1*μ*M serotonin on the pyrolytic electrode and a pyrolytic electrode coated with CNTs. **A**: Graphite electrodes show the oxidation and reduction of serotonin at 400 V s^-1^.0.7V (±0.1V) and reduced at −0.1V. **B:** Shows the oxidation of serotonin on pyrolytic-CNT electrodes at scan speed 600V s^-1^. 0.65 ±0.18V and reduction peak at 0.2 V±0.1V. **C:** Shows the application of “Serotonin Waveform” (Hashemi et al., 2009; Jackson et al., 1995) wherein, multistep oxidation of serotonin can be seen at 0.72V(±0.05V) and a secondary oxidation peak is seen at 0.45V, a reduction peak can be seen at 1.1V and −0.3V. **D**: Shows the plot of scan rate *vs* oxidation peak current. The electrodes, having a mean diameter of 5μm and a nominal height of 0.9μm, show the sensitivity of 5±17nA/μM. Data is normalized. **E**: Simulations of oxidation of serotonin where the diffusion coefficient is 5 X 10^-6^ cm^2^/m, alpha is set to 0.4, the number of electron transfer (n)=1 and the rate of electron transfer is 3 X 10^-6^. It should be noted that upon increasing the number of electron transfers to 2 a reduction peak can be obtained **F**: Shows a 2D-hotspot, wherein the voltage is at X-axis, repetition is at Y-axis, and current at Y’-axis. Multiple oxidation and reduction states can be seen at −0.3V and 1.3V, while broad oxidation peaks can be seen at 0.7V, which corresponds to the oxidation of serotonin.

### 2. Selectivity and Specificity

The *in vivo* environment contains multiple analytes which tend to interfere with the determination of the analyte of interest. These analytes cause non-specific adsorption on the surface of the electrode which limits the sensitivity of the target molecule and the probing time (Swamy and Venton, 2007). Dopamine is a common interference that is encountered in the striatum because the striatum receives dense and direct projections from the ventral tegmental area (VTA) and substantia nigra (SN) (Schubert et al., 2015). To understand the dopamine response for serotonin waveform, 1*μ*M of dopamine was introduced to the flow cell, and “Serotonin Waveform” with modifications was applied (Hashemi et al., 2009; Jackson et al., 1995). Dopamine showed an incomplete oxidation peak at 0.4V (**Figure 2A**) The sensitivity for dopamine, using the serotonin waveform was 52nA/*μ*M, thus showing that the waveform is not suitable for dopamine detection.

**Figure 2:**
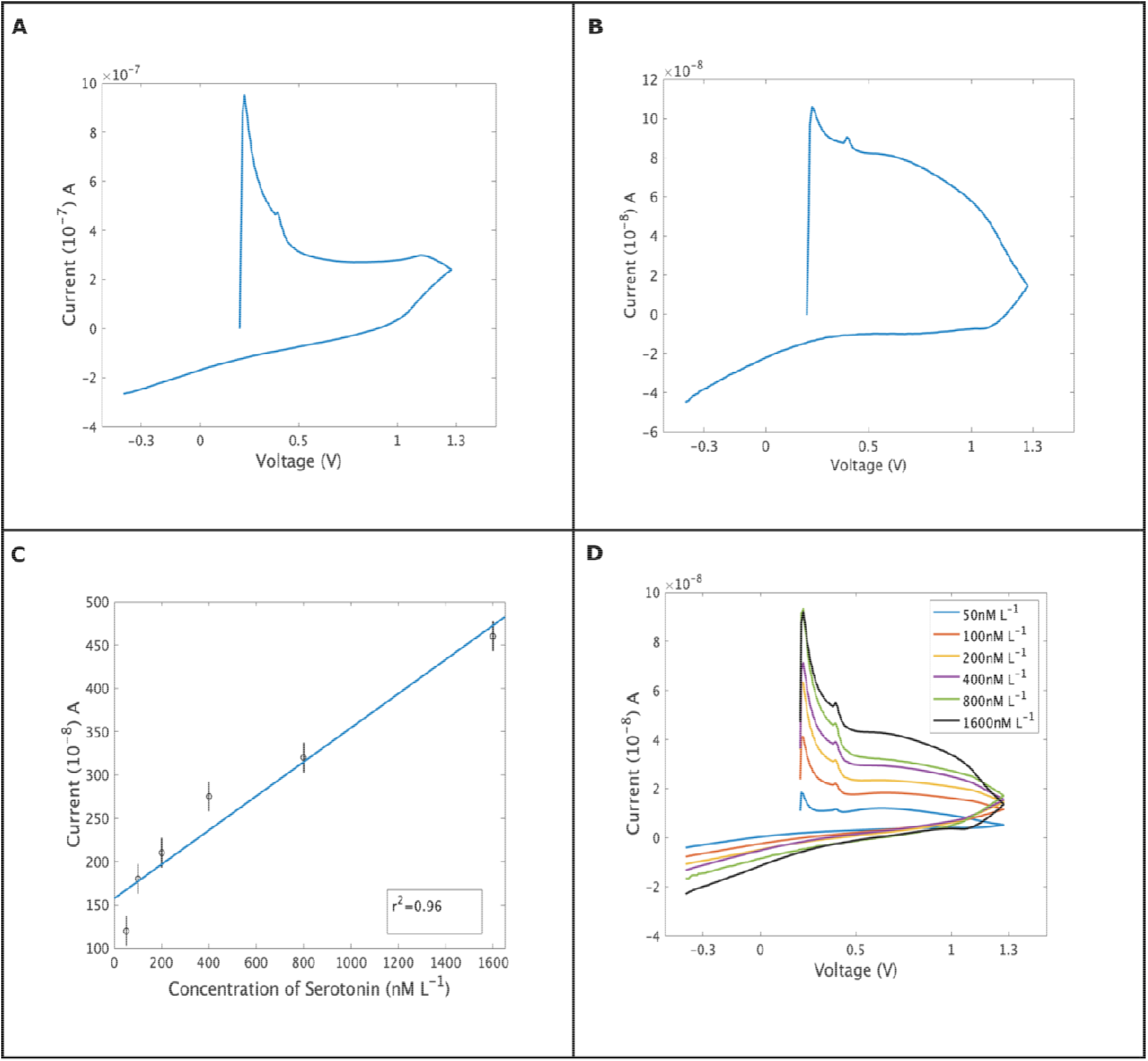
Shows the selectivity of pyrolytic-CNT coated electrodes. **A:** 1*μ*M of dopamine using serotonin waveform. An incomplete oxidation peak is seen at 0.4V. **B:** Electrochemical response to 10*μ*M of 5-HIAA. An oxidation peak can be seen at 0.85V. **C:** Calibration curve for serotonin. The graph has a fit of r^2^=0.96. **D:** Trials against concentrations.

Electrochemical detection of serotonin is confounded by 5-Hydroxyindoleacetic acid (5-HIAA), which is an oxidation product of serotonin (Hashemi et al., 2009; Swamy and Venton, 2007). The high concentration of 5-HIAA results in adsorption of 5-HIAA on the surface of the electrode, thereby slowing the temporal kinetics of electrodes (Swamy and Venton, 2007). We thus measured 5-HIAA, by injecting 10μM in 6 cycles in a flow cell apparatus using the modified serotonin waveform. The results showed that 5-HIAA oxidizes at 0.85V (**Figure 2B**), distinct from serotonin oxidation at 0.2V. Finally, the electrode was exposed to known concentrations of serotonin from 50 to 1600 nM (**Figure 2C and Figure 2D**). The sensor showed linearity in this range and had a detection limit of 20 ±9nM/L.

### 3. MultiModal Sensing

Dopamine and serotonin are found in the striatum wherein they both are co-expressed abundantly. In order to study the multimodal detection of dopamine and serotonin simultaneously, we first used pyrolytic electrodes for distinguishing between dopamine and serotonin. Uncoated pyrolytic electrodes were unable to spatially resolve dopamine and serotonin and showed only 1 oxidation and reduction peak at 0.5V and 0.0V (**Figure 3A**). After CNT coating, we noticed 2 clear oxidation peaks, 0.1V for dopamine 0.5V for serotonin and reduction for serotonin is seen at 0.1V, and dopamine is reduced at −0.3V (**Figure 3B**) using a voltage triangular waveform, i.e −0.4V to 1.3V and cycled back at −0.4V at 1000V s^-1^. Although a previous study had previously shown simultaneous detection of dopamine and serotonin (Swamy and Venton, 2007), due to the single oxidation peak for both dopamine and serotonin, thus far it was not possible to detect the two analytes independently and simultaneously. In our current study, our base material was pyrolyzed carbon was coated with CNTs which had heteroatoms, which allowed us to distinguish oxidation and reduction peaks.

**Figure 3.**
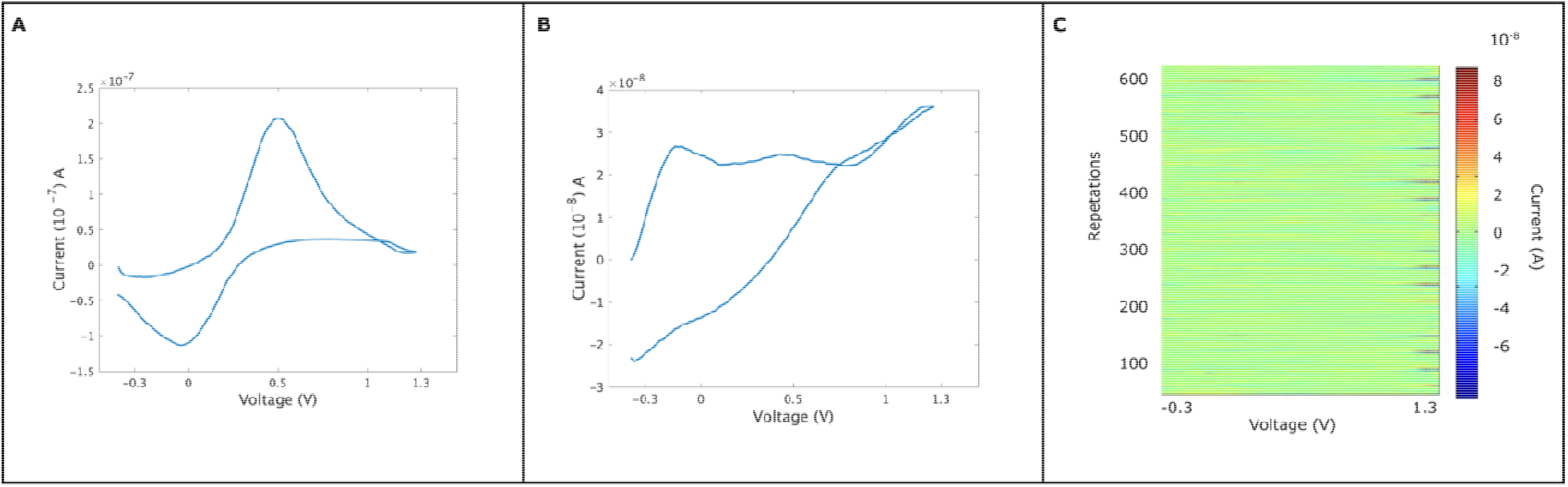
Cyclic Voltammogram for simultaneous detection of dopamine and serotonin using a pyrolytic electrode and pyrolytic-CNT electrode. **A:** Shows the oxidation of 1*μ*M dopamine and serotonin simultaneously on pyrolytic electrodes. One single oxidation peak can be seen at 0.58V and a reduction peak at 0.0V can be seen. **B**: Shows the oxidation for dopamine at 0.1V and serotonin at 0.5V. Serotonin is reduced at 0.1V and dopamine is reduced at −0.3V. **C:** 2-D hotspot wherein multiple oxidation and reduction states can be observed.

### 4. Mechanism of Serotonin Oxidation of Pyrolytic-CNT electrode

To investigate the redox dynamics of serotonin oxidation on pyrolytic electrodes and pyrolytic-CNT electrodes, we performed simulations using DFT. The interactions of serotonin with graphite surfaces occur, wherein graphite surfaces are disrupted (**Figure 4A and Figure 4B**). This remains consistent with our experimental observations wherein serotonin fouls the graphite surfaces, thus limiting the shelf life of graphite-based probes. While CNT and serotonin simulations show that the binding of serotonin to CNT is not favorable. The initial bond distance between serotonin and CNTs was (8.4□) while as the simulation progress the bond distance between serotonin and CNTs drops to (5□) followed by an increase in the bond distance again to (8□). This suggests that CNT surfaces do not allow the bonding of serotonin molecules. Even when surfaces are functionalized with the oxygen group (Figure 4C and 4D) the interaction is limited to (4□), thus suggesting that CNTs surfaces limit the interaction of serotonin molecules. These observations remain consistent with our experimental data, thus suggesting that surface roughness, texture and chemistry plays an important role in dictating the electrochemical response of serotonin.

**Figure 4.**
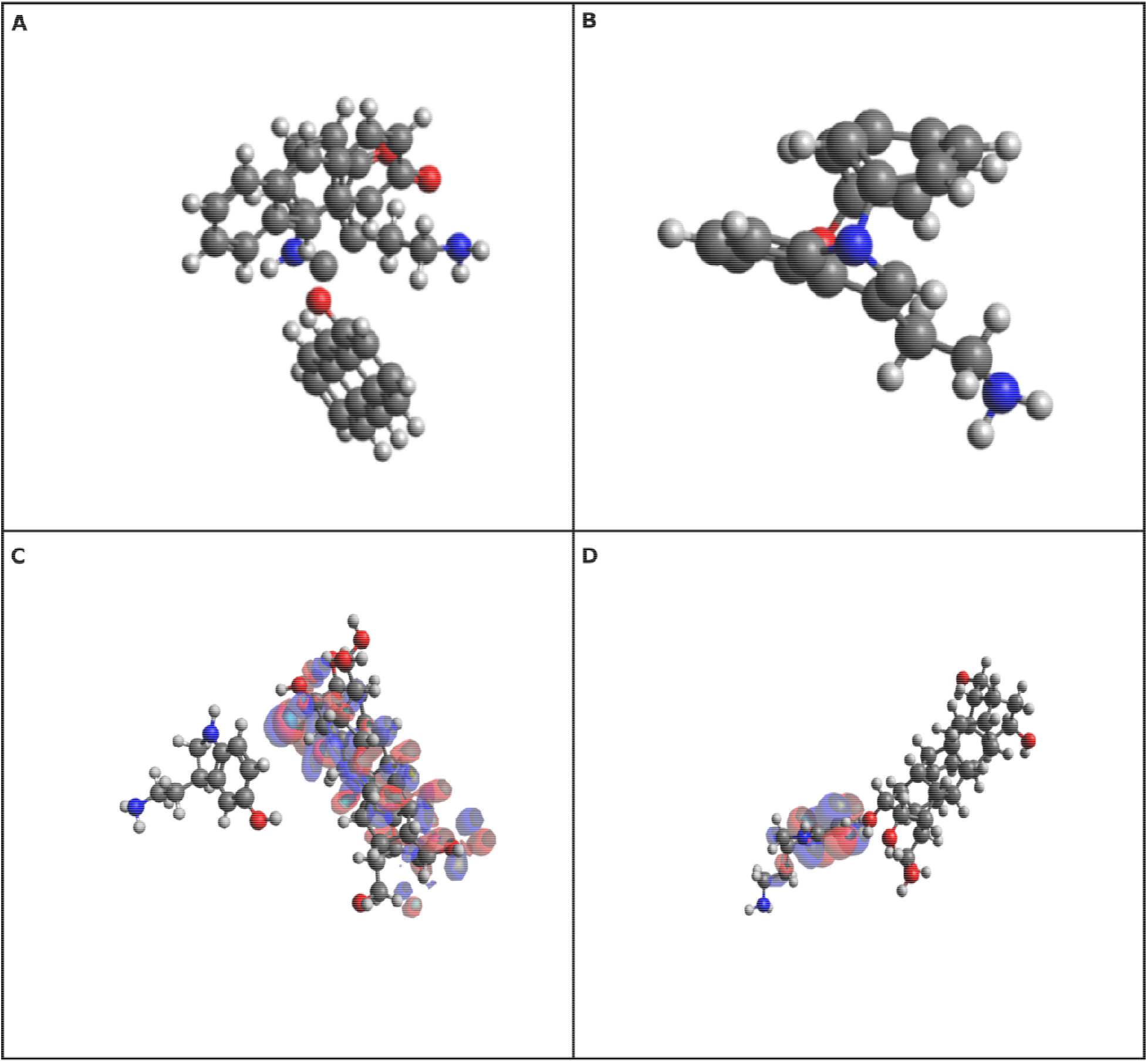
Serotonin oxidation on carbonized surfaces. **A:** Shows the disruption of the graphite surface when serotonin is introduced to the system. The graphite surface loses its pi-pi stacking and forms a bond with the serotonin molecule. **B:** Binding of serotonin to monolayer carbon surface (graphene). Serotonin forms carbon-carbon single bonds with the nitrogen group on the graphene surface. C: The highest occupied molecular orbitals (HOMO) for serotonin, using chains functionalized with hydrogens. It can be seen that hydrogenated surfaces limit the interaction of serotonin. **D:** Shows the interactions of oxygen functionalized CNT surfaces. The interactions of serotonin with oxygen motifs are higher than hydrogenated surfaces, however, no binding occurs, thus showing the antifouling properties of CNT surfaces.

## Conclusions

In this work we showed that graphite-based surfaces can detect serotonin, the occurrence of biofouling limits the shelf life of graphite-based surfaces. Upon adapting with CNT coating, we found an increase in sensitivity, selectivity, and the possibility of multimodal detection of dopamine and serotonin. Our future work will be *in vivo* detection of serotonin and dopamine simultaneously.

## Acknowledgements

We would also like to thank Geert-Jan Janssen for his help with electron microscopy. This work was supported in part by European Regional Development Fund (MIND, nr. 122035).

## Supplementary Information-I

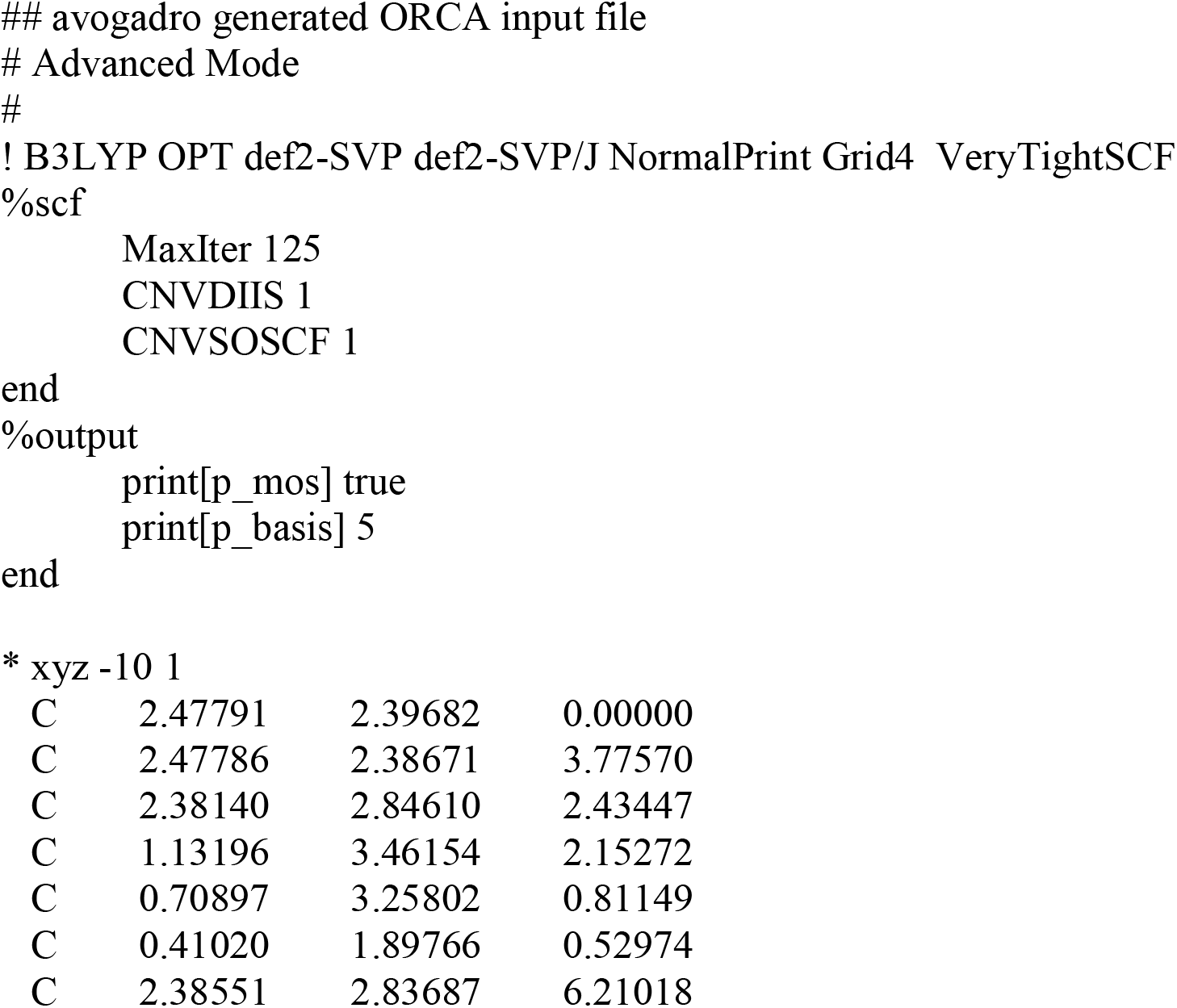
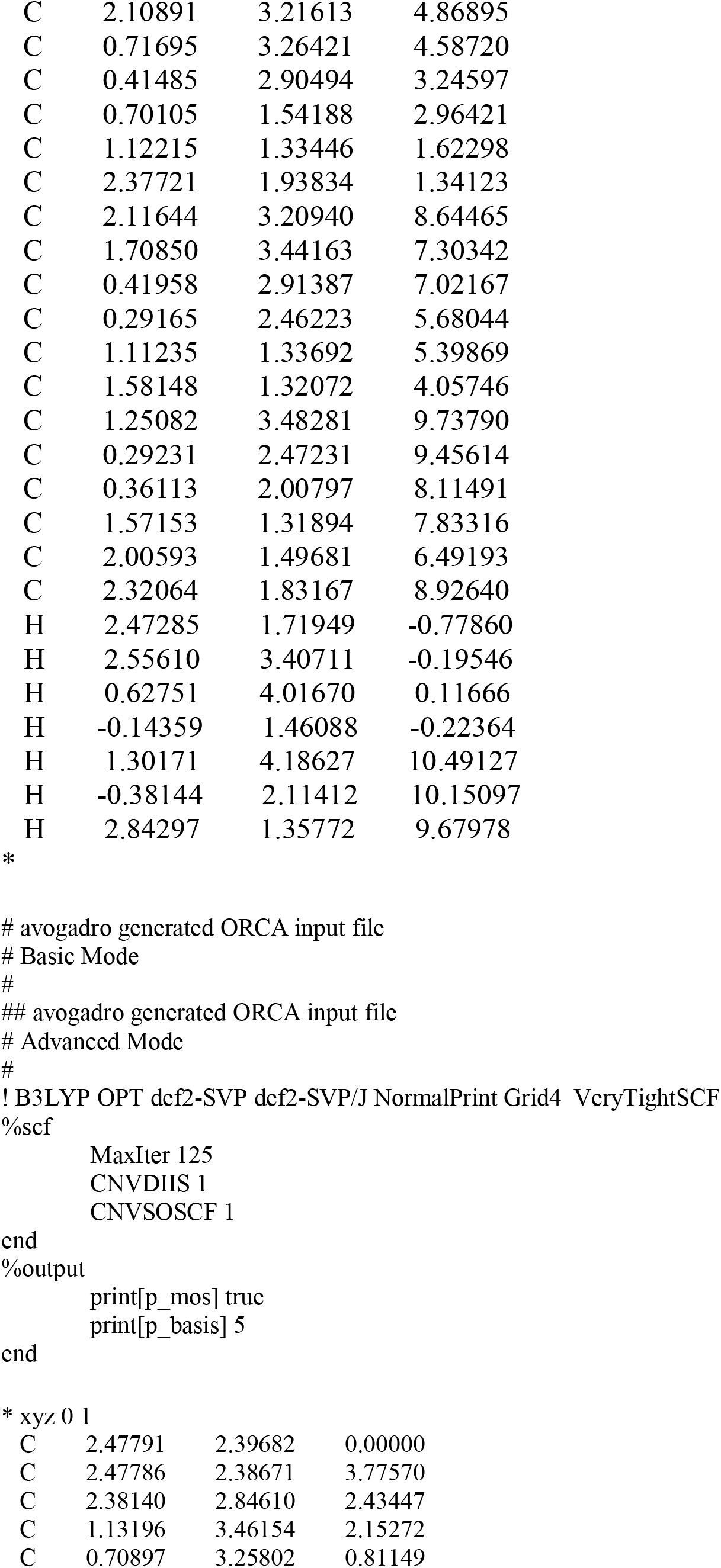
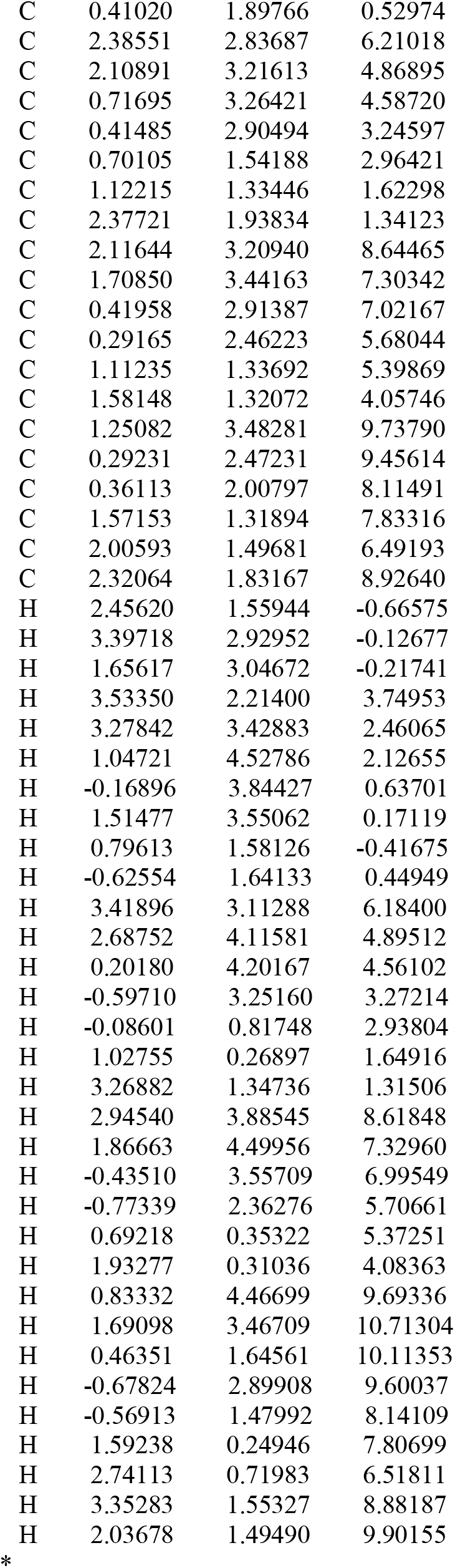

## Supplementary Information-II

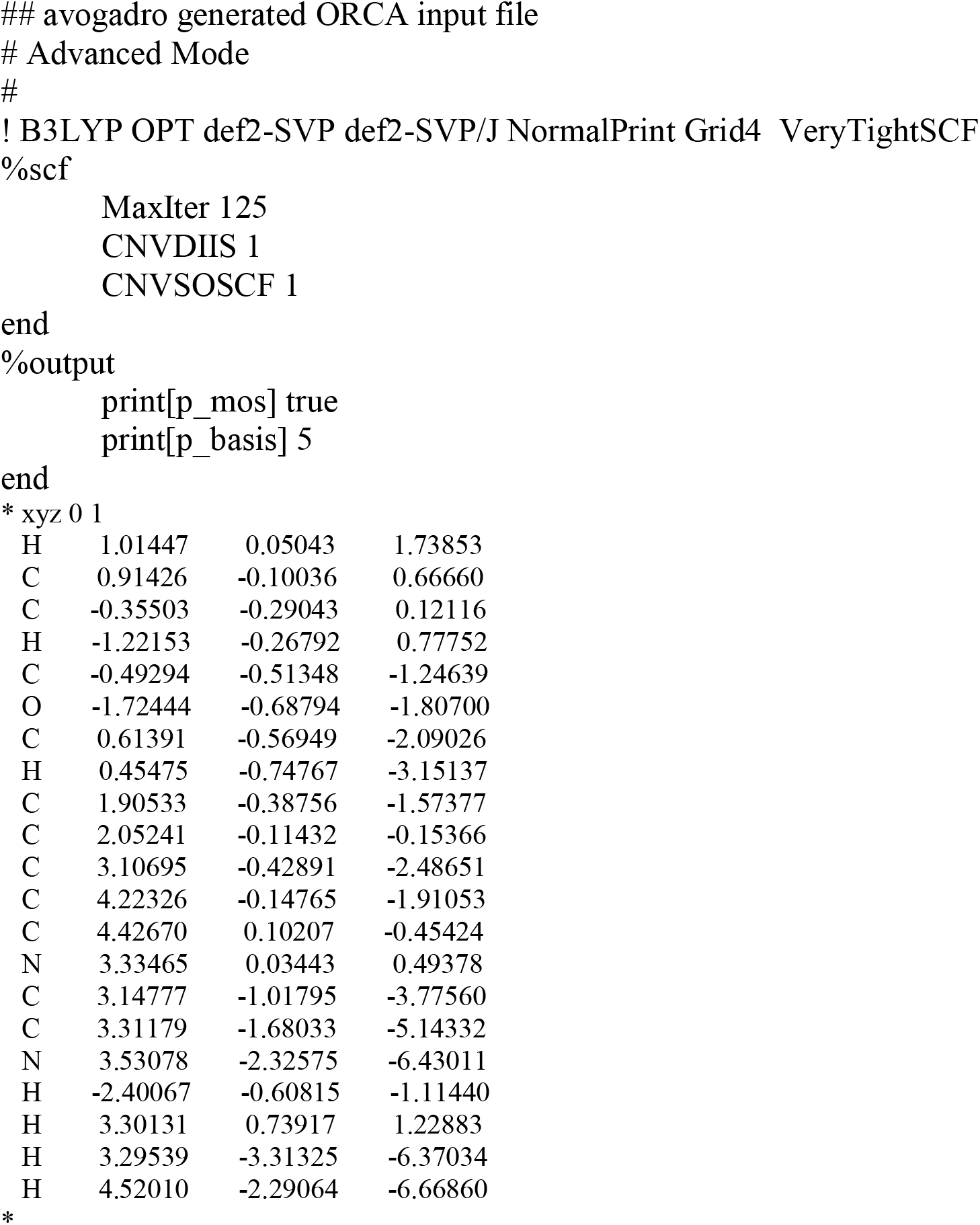

## Supplementary Information-III

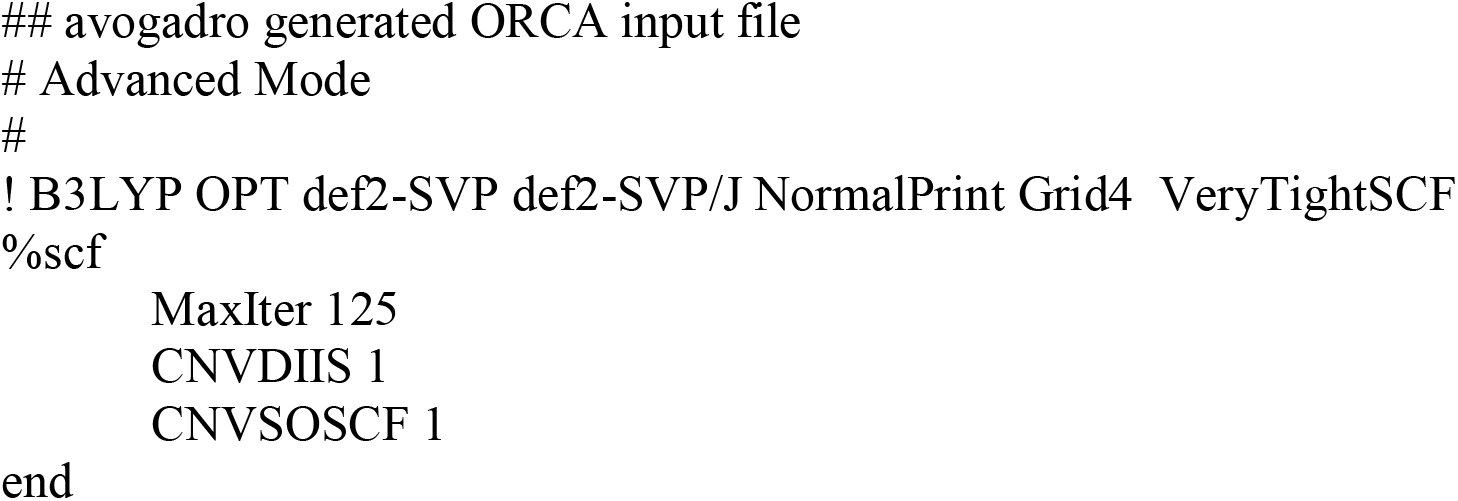
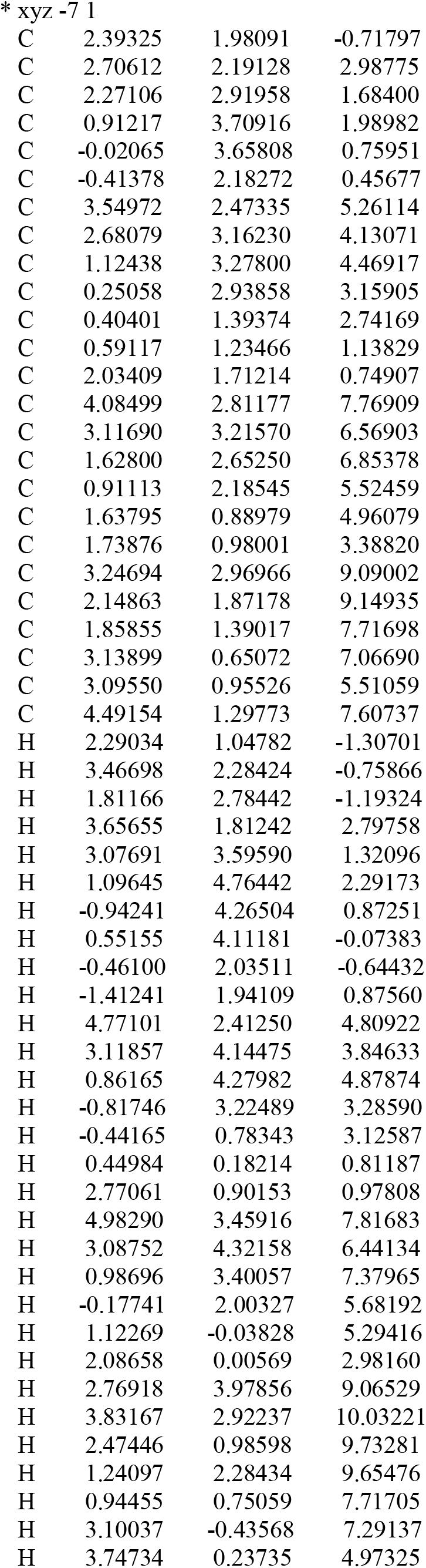
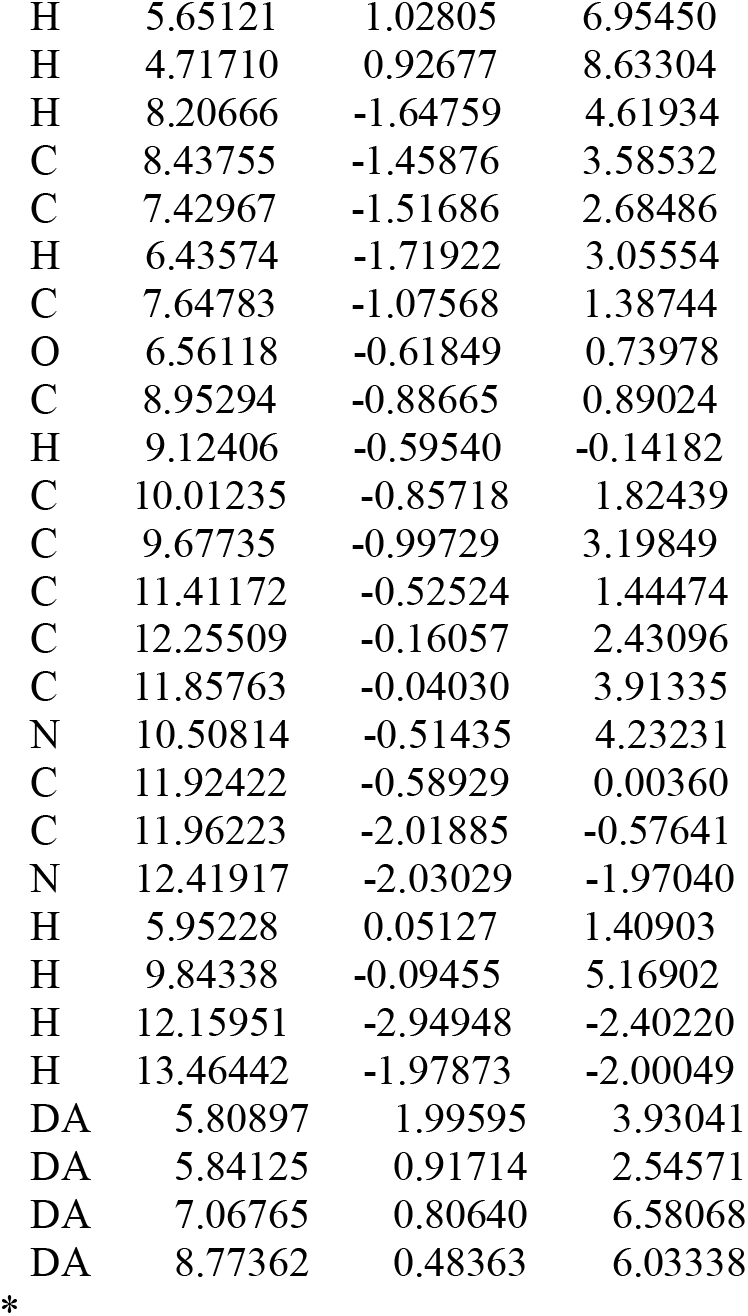

## Notes

### Competing Interest Statement

The authors have declared no competing interest.

### Summary of Updates

The title and order of authors has been changed to be more consistent and to make the paper easier to find. The rest of the text is unchanged

